# A Two-tiered Functional Screen Identifies Herpesviral Transcriptional Modifiers and their Essential Domains

**DOI:** 10.1101/2021.09.20.460982

**Authors:** David W Morgens, Divya Nandakumar, Allison L Didychuk, Kevin J Yang, Britt Glaunsinger

## Abstract

While traditional methods for studying large DNA viruses allow the creation of individual mutants, CRISPR/Cas9 can be used to rapidly create thousands of mutant dsDNA viruses in parallel. Here, we used this approach to study the human oncogenic Kaposi’s sarcoma-associated herpesvirus (KSHV). We designed a sgRNA library containing all possible ~22,000 guides targeting the genome of KSHV – one cut site approximately every 8 base pairs – enabling the pooled screening of the entire genome. We used this tool to phenotype all possible Cas9-targeted viruses for transcription of KSHV late genes, which is required to produce structural components of the viral capsid. By performing targeted deep sequencing of the viral genome to distinguish between knock-out and in-frame alleles created by Cas9, we discovered a novel hit, ORF46 – and more specifically its DNA binding domain – is required for viral DNA replication. Our pooled Cas9 tiling screen followed by targeted deep viral sequencing represents a two-tiered screening paradigm that may be widely applicable to dsDNA viruses.

## Introduction

Human oncogenic viruses are a major cause of cancer, with recent estimates that 15% of all cancers are associated with a viral infection (Zapatka *et al.*, 2020). Gammaherpesviruses are a class of double-stranded DNA viruses that include both the oncogenic Epstein-Barr virus (EBV) – a ubiquitous infection associated with a number of malignancies (Thompson and Kurzrock, 2004) – and Kaposi’s sarcoma-associated herpesvirus (KSHV), a major cause of cancer in AIDS and other immunocompromised patients (Ganem, 2010; Minhas and Wood, 2014). While many of the viral factors involved in replication and transcription of the KSHV genome have been identified, numerous non-coding regions, transcripts, and ORFs have no ascribable function (Toth *et al.*, 2010; Arias *et al.*, 2014; Ye, Zhaoid and Karijolich, 2019; Brackett *et al.*, 2021).

During the lytic stage of KSHV replication, the virus expresses its genes in kinetically regulated waves, culminating in the formation and release of infectious virions. These include immediate early genes that regulate reactivation of the virus, early genes that encode components required for viral DNA replication, and finally late genes, which encode for capsid proteins required for virion formation. Immediate early and early genes are driven by host-like promoters, while late gene transcription in KSHV and related gammaherpesviruses has several unique features including truncated promoters and modified TATA boxes (Serio *et al.*, 1998; Tang, Yamanegi and Zheng, 2004; Wong-Ho *et al.*, 2014; Davis *et al.*, 2015; Gruffat, Marchione and Manet, 2016). They also require at least six virally encoded transcriptional activators (vTA) (Aubry *et al.*, 2014; Davis *et al.*, 2016) and, perhaps most notably, their transcription is dependent on viral DNA replication (Wang *et al.*, 2008; Aubry *et al.*, 2014; Djavadian, Chiu and Johannsen, 2016).

The genomes of herpesviruses are double-stranded DNA that replicate in the nucleus and are thus targetable by CRISPR/Cas9. Indeed, CRISPR-based targeting of herpesviruses has been explored as a way to suppress infection (Wang and Quake, 2014; Roehm *et al.*, 2016; van Diemen *et al.*, 2016; Karpov *et al.*, 2019; Tso, West and Wood, 2019; Walter and Verdin, 2020). CRISPR has also been used to make functional insertions and knockouts on the viral genome (Suenaga *et al.*, 2014; Yuen *et al.*, 2015; BeltCappellino *et al.*, 2019) or to perform pooled functional screens (Hein and Weissman, 2019; Gabaev *et al.*, 2020) in much the same way as on the mammalian genome. The use of CRISPR to identify essential domains of proteins (Shi *et al.*, 2015; He *et al.*, 2019) has yet to be applied to dsDNA viruses, but has particular potential given that viral proteins are frequently multifunctional.

Here, we applied a CRISPR approach to probe for functional regulators of KSHV late gene expression by creating a library of sgRNAs that tiles the KSHV genome, corresponding to one sgRNA for every 8 base pairs. Using this tiling library, we systematically screened for regulators of viral late gene transcription. In addition to capturing the majority of known components of this system, we identify novel viral regulators of late gene expression along with higher resolution data highlighting essential domains of these proteins. We describe a novel role for KSHV ORF46 in late gene transcription through its requirement for KSHV DNA replication. Targeted deep sequencing of the ORF46 locus provided additional mechanistic insight by identifying the mutations created by Cas9 and implicating an essential role for its DNA binding but not its catalytic domain, which we confirmed through reverse genetics experiments. Collectively, this pipeline provides a framework to create and phenotype thousands of viral mutants to gain high-resolution functional information on KSHV and other dsDNA viral genomes.

## Results

### Pooled tiling screen of KSHV genome for modifiers of late gene expression

To identify viral modifiers of late gene expression, we developed a Cas9+ reporter system. We first infected the human renal carcinoma cell line iSLK (Myoung and Ganem, 2011; Brulois *et al.*, 2012) with a modified version of the BAC16 KSHV genome containing a far-red fluorescent protein driven by the promoter of the viral late gene K8.1 as well as a constitutive EGFP (**Figure S1a**). Upon reactivation by doxycycline-induced ORF50 – the major lytic transactivator – and sodium butyrate, this cell line maintained viral activity comparable to wildtype and that the reporter recapitulated key features of late gene expression after reactivation, including expression 48 hours post reactivation and sensitivity to inhibitors of viral DNA replication (**Figure S1b-d**). To confirm our ability to create functional mutants, we lentivirally delivered Cas9 along with sgRNAs targeting various regions of the KSHV genome into a BAC16-infected iSLK line and used a supernatant transfer assay to evaluate their impact on virion production (**Figure S2a**). Although the guides targeting the essential viral loci elicited the strongest reduction in infectious virion production, we note that sgRNAs targeting some nonessential regions of the viral genome did appear to have a functional effect above background, perhaps through DNA damage on the viral genome (**Figure S2b**). By delivering Cas9 to the iSLK line infected with the reporter virus, we can measure the effect of viral-targeting guide RNAs on the expression of late genes.

We then used this Cas9+, late gene reporter line to screen an sgRNA library containing all possible ~22,000 sgRNAs targeting the viral genome (**Supplementary Data 1**). After lentiviral delivery of the library, cells were chemically induced to enter the lytic cycle through activation of the ORF50 major lytic transactivator and sorted by FACS based on their reporter expression level. Composition of high and low late-gene expression fractions were quantified by next-generation sequencing of the sgRNA locus (**Figure 1a; Supplementary Data 2**). Enrichment of each guide was reproducible between two replicates, representing a phenotype every ~8 bp across the KSHV genome (**Figure S3a,b**).

**Figure 1.**
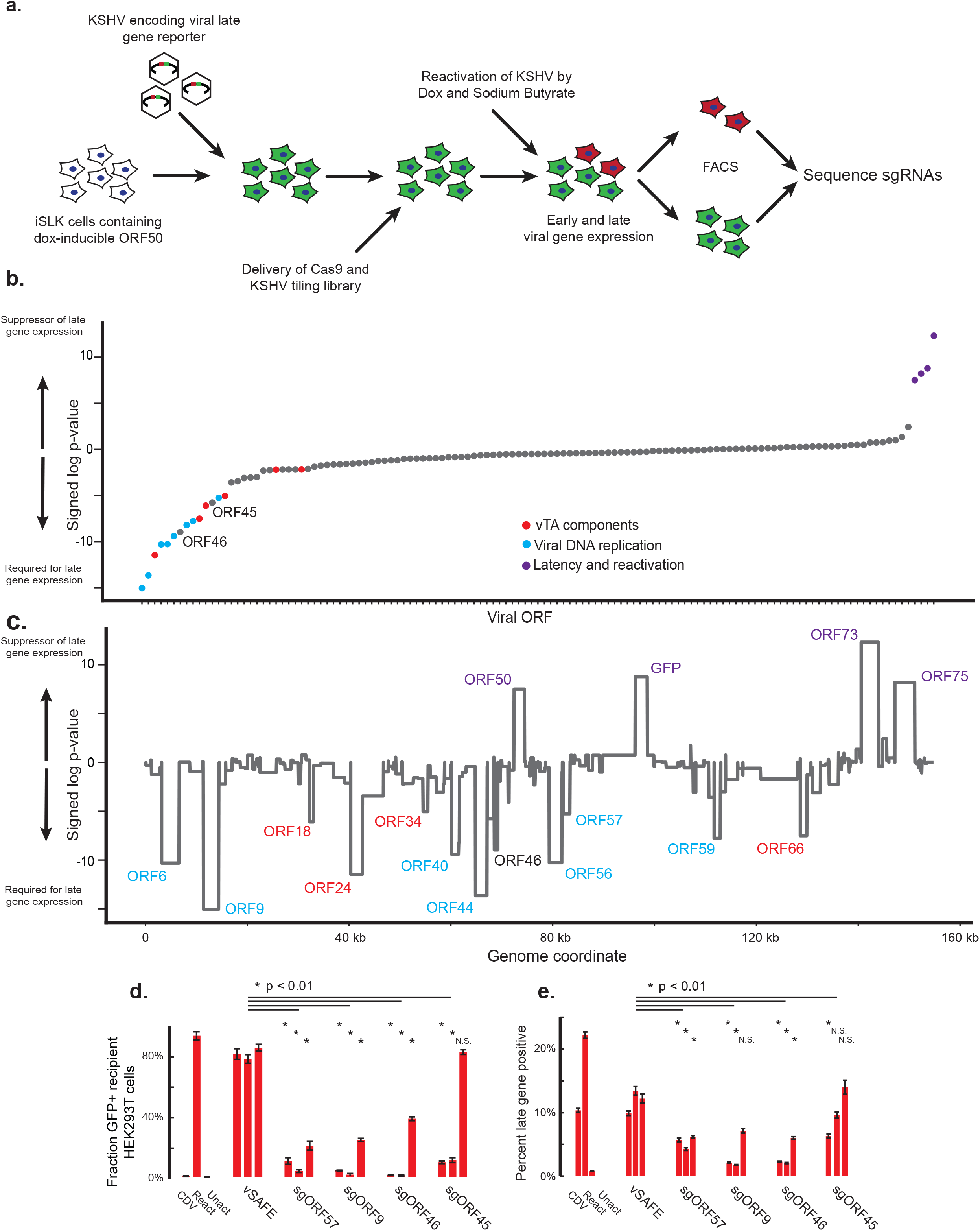
Pooled tiling screen of KSHV genome for modifiers of late gene expression. a) Design of viral tiling screen for late gene expression. iSLK cells were infected with KSHV encoding a far-red reporter of late-gene activity. Cells were lentivirally infected with Cas9-blast and subsequently the KSHV tiling sgRNA library. After 1-2 weeks of latency to allow for editing, cells were reactivated into the lytic cycle by doxycycline and sodium butyrate. 48 hours post reactivation, cells were trypsinized, fixed, and sorted for high and low late-gene expression. The sgRNA locus in each population was then sequenced to calculate enrichment. b,c) To calculate significance, sgRNAs targeting each annotated region of the genome were grouped and a signed log Mann-Whitney p-value was calculated comparing each viral region to the negative control sgRNAs using an average enrichment from two replicates. Viral ORFs sorted by significance (b) or genome location (c). A positive log p-value indicates this region promotes late gene expression when disrupted, and a negative log p-value indicates this region is required for expression of late genes. d,e) Three independent sgRNAs from the screen targeting each indicated ORF were individually cloned and delivered to late-gene reporter iSLK cells. Viral activity was measured by supernatant transfer to uninfected HEK293T cells (d) and late-gene reporter activity (e). Error bars are standard error from three independent reactivations. Activated, unreactivated, and parent samples treated with the viral DNA replication inhibitor cidofovir (CDV) were included as controls, along with three vSAFE guides targeting an unexpressed region of the viral BAC. P values were calculated using a single-tailed, equal variance Student’s t-test, with the least significant value used when compared to each individual vSAFE sgRNA. NS indicates non-significance (P>0.01).

By comparing the distribution of sgRNA phenotypes targeting each KSHV ORF to the background of negative control sgRNAs, we reproducibly identified many ORFs strongly required for late-gene expression (**Figure 1b,c; Figure S3c; Supplementary Data 3,4**). These include all six known components of the viral DNA replication machinery (ORF6, ORF9, ORF40/41, ORF44, ORF56, and ORF59) along with 4 of the 6 vTA components (ORF18, ORF24, ORF34, and ORF66). In addition, we identified ORF57, which is known to be required for the efficient expression of ORF6 and other DNA replication components (Verma *et al.*, 2015). A viral tegument protein, ORF45 (Zhu and Yuan, 2003), showed mixed results in validation and was not pursued for further characterization (**Figure 1d,e**). ORF46, a DNA repair enzyme (Earl *et al.*, 2018) not previously associated with late gene transcription, also showed severe defects in late gene expression which we subsequently validated with individual guides (**Figure 1d,e**). Together, this demonstrates the identification of both known and novel viral modifiers of late gene expression.

### Deep viral sequencing demonstrates ORF46 and its DNA binding activity are required for viral DNA replication

To further investigate the role of the novel regulator ORF46, we deep sequenced several viral loci to identify and phenotype mutations created by Cas9 nuclease activity (**Figure 2a**). This approach enables us (1) to distinguish between Cas9-created mutations that result in a knockout versus an amino acid change and (2) to screen for functional defects associated with a given mutation using phenotypes such as viral DNA replication and virion production (**Figure S4a**). We amplified and sequenced ORF46 – along with one negative control locus, ORF21, and two components of the vTA complex, ORF18 and ORF34 – from cells in three different viral states. These included (1) latently infected cells (where mutations should have little effect), (2) cells 48 hours post reactivation, at which time the viral genome is being replicated, and (3) from supernatant 72 hours post reactivation when virions are released into the media (**Figure 2a; Supplementary Data 5; Supplementary Table 1**). We first identified out-of-frame mutations and observed a depletion in ORF46 knockout alleles in cells undergoing viral DNA replication as well as a depletion of knockout alleles from viral DNA in the supernatant (**Figure 2b**). In contrast, knockout alleles of vTA components ORF34 and ORF18 were only depleted in supernatant samples, consistent with their specific requirement after viral DNA replication (Figure 2b; Figure S4b). Knockout mutations in a nonessential control, ORF21, are mildly depleted in both samples (**Figure 2b, Figure S4b**); this is consistent with the previously observed effect of Cas9 activity on viral activity (**Figure S2b**). The relative depletion of ORF46 knockouts from replicating viral DNA suggests that ORF46 acts prior to late gene expression and is required for viral DNA replication.

**Figure 2.**
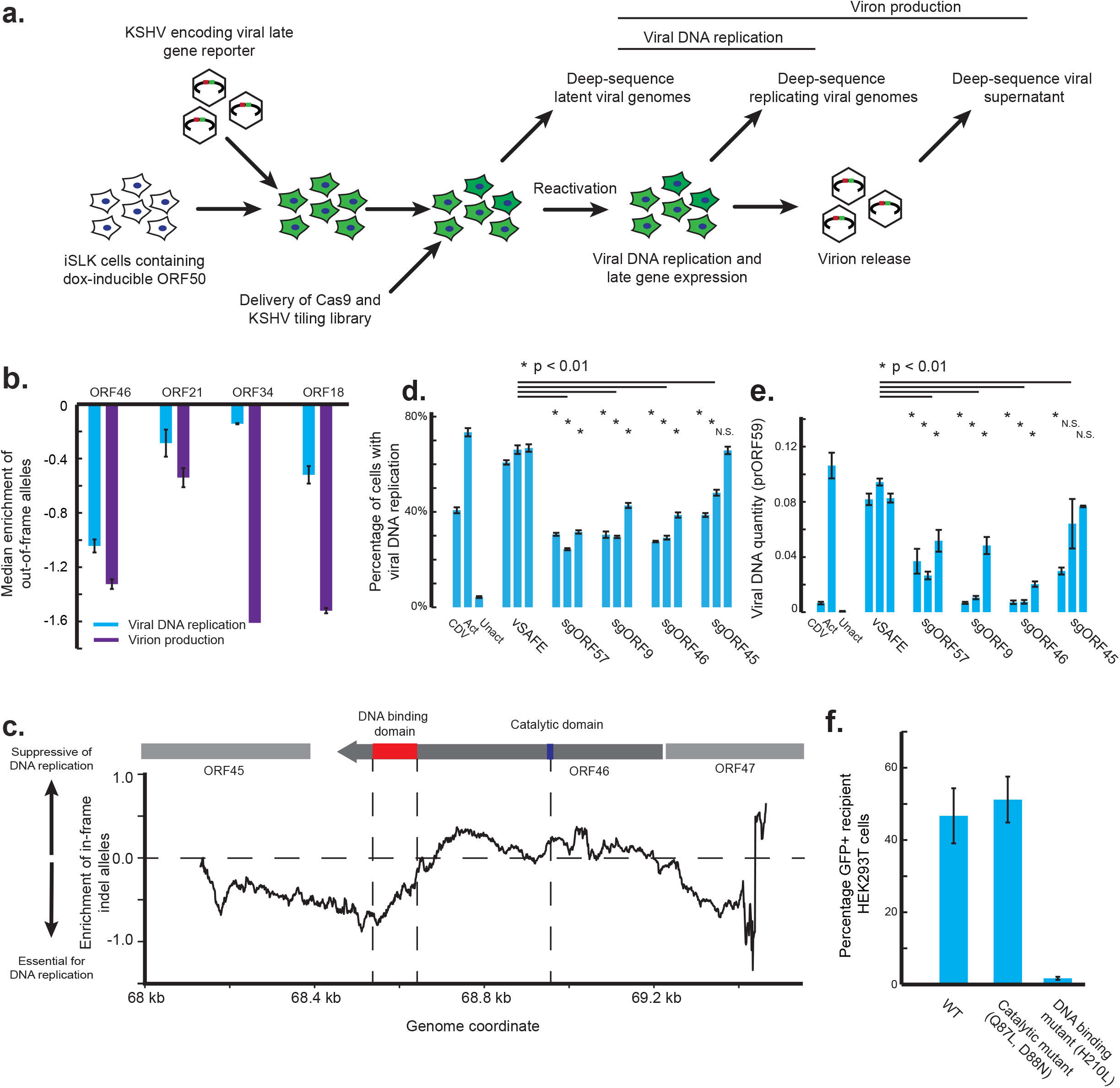
Targeted deep sequencing reveals essential domains of ORF46. a) Design of targeted deep-sequencing experiment. Four viral loci were amplified and deep-sequenced: ORF21, ORF18, ORF34, and ORF46. b) Median enrichment across the coding region of the gene of out-of-frame indels in replicating cells relative to latent cells (Viral DNA replication) or the supernatant relative to latent cells (Virion production). Error bars are standard error from two replicates. One replicate of ORF34 supernatant sample was excluded due to uneven coverage. c) Smoothed signal from in-frame mutations across the ORF46 loci in comparison between latent and replicating genomes. In-frame mutations were defined as insertions or deletions whose size where divisible by three. The number of mutations was pooled in a 100 bp window and enrichment was calculated relative to the enrichment of out-of-frame mutations. A negative value indicates that in-frame mutations were relatively depleted at the indicated region. d,e) Three individual sgRNAs were lentivirally delivered to KSHV-infected iSLK cells targeting the indicated viral ORF. Error bars are standard error from four independent reactivations. CDV-treated, reactivated, and unactivated parental cells are included as controls, along with three vSAFE sgRNAs targeting an ORF-free region of the BAC. P values were calculated using a single-tailed, equal variance Student’s t-test, with the least significant value used when compared to each individual vSAFE sgRNA. NS indicates non-significance (P>0.01). d) Measuring viral DNA replication by percentage positive EdU staining. A 2-hour pulse of EdU was delivered 48 hours post reactivation. Cells were then fixed, and click chemistry was used to measure EdU incorporation by flow cytometry. Unreactivated cells were used to establish gates for flow cytometry, where subgenomic EdU incorporation was measured as viral DNA replication. (e) DNA was extracted 48 hours post reactivation, and qPCR of the viral promoter of ORF59 was used to measure the amount of viral DNA. f) Viral activity was measured for iSLK cells infected with KSHV encoding either wildtype ORF46, a catalytic mutant of ORF46 (Q87L, D88N), or a DNA-binding mutant of ORF46 (H210L). Cells were reactivated, and after 72 hours supernatant was filtered and transferred to uninfected HEK293T cells. Infection was monitored by expression of the BAC16-encoded, constitutive EGFP. Error bars are standard error from three technical replicates.

By identifying in-frame mutations, we can further refine what regions of these proteins are essential for their role in the viral life cycle, analogous to previous work on the human genome(Shi *et al.*, 2015). Across each of the loci, we examined the relative depletion of in-frame alleles between replicating DNA and supernatant for ORF18 and ORF34 – and between latent virus and replicating DNA for ORF46. Indeed, depletion of in-frame alleles in ORF18 corresponded with previously identified residues required for interaction with other vTA complex members ORF30, ORF31, and ORF66 (**Figure S4c; Supplementary Data 6**) (Castañeda and Glaunsinger, 2019), and several regions observed in ORF34 correspond to previously phenotyped truncation mutants with defects in binding the vTAs ORF24 or ORF66 (**Figure S4d**) (Nishimura *et al.*, 2017). In ORF46, we noted that these in-frame mutations near the C-terminus were depleted from replicating DNA samples relative to the rest of the coding region (**Figure 2c**). This suggests that the DNA-binding domain encoded in the C-terminus is required for viral DNA replication but that the catalytic domain is not. We also observed signal outside the coding region of ORF46, which may correspond to disruption of the adjacent ORF45 or perturbation to upstream regulatory elements of ORF46.

We next tested our predictions that ORF46 and more specifically its DNA binding activity is required for viral DNA replication using orthogonal methods. First, we delivered the panel of sgRNAs from Figure 1d into an independent KSHV-infected Cas9+ iSLK line; the sgRNA panel included targets for both ORF57 and ORF9, both of which are required for viral DNA replication, as well as for ORF46 and ORF45. Guides targeting ORF46 interfered with incorporation of EdU, suggesting a viral DNA replication defect (**Figure 2d**), which was confirmed for the ORF46 sgRNAs by qPCR of viral DNA (**Figure 2e**). Guides targeting ORF45 showed inconclusive effects. Notably, none of the genes targeted in the above panel, including ORF46, was required for early gene expression, as measured by expression of the HaloTag-fused ORF68 early gene (**Figure S4e**). Together, these data support a specific requirement of ORF46 for viral DNA replication, which is bolstered by data with the ORF46 homolog in EBV (Su *et al.*, 2014). Notably, ORF46 disruption phenocopied disruption of the viral DNA polymerase ORF9, demonstrating the key role ORF46 plays in viral DNA replication. To test more specifically whether this activity is dependent on the DNA binding activity of ORF46, we individually mutated residues required for DNA binding (H210L) or catalysis (Q87L, D88N) in ORF46 using BAC mutagenesis (Su *et al.*, 2014). Reactivation of these mutant viruses showed that the DNA binding domain but not the catalytic domain of ORF46 is required for the viral life cycle (**Figure 4f**). Together these data orthogonally confirm our prediction that ORF46 and its DNA binding domain are required for viral DNA replication.

## Discussion

Late gene expression in oncogenic gammaherpesviruses is a mechanistically unique, essential and potentially targetable viral process. Here we applied an exhaustive tiling approach to KSHV to identify viral proteins required for expression of viral late genes. We identified a large majority of the known late gene regulators and confirmed that one additional protein, ORF46, is required for this process (**Figure 1**). Targeted deep sequencing of the viral mutants created by Cas9 suggested that the DNA binding domain of ORF46 is required for viral DNA replication, a prediction we validated through viral mutagenesis (**Figure 2**). This two-level screening approach is broadly applicable to connecting phenotype to genotype for a variety of processes involved in the KSHV lifecycle – as well as more broadly for other dsDNA viruses.

Compact viral genomes present both unique challenges and opportunities for genomics-based approaches to study viral phenotypes. On one hand, the high functional density in a relatively small genome allows approaches such as ours to perturb every element. For example, here we tile all ~80 ORFs and their cis regions with fewer guides than previous studies have used to tile just three human genes (Tycko *et al.*, 2019). On the other hand, viral loci often encode multifunctional proteins and can also encompass overlapping ORFs and noncoding regulatory sequences, which means that some mutations may impact multiple viral features(Arias *et al.*, 2014). This is true for our study as well, as the ORF46 DNA binding domain also overlaps with the promoter and TSS of the adjacent gene ORF45. Thus, although the CRISPR nuclease is less likely to perturb non-coding elements (Tycko *et al.*, 2019), some of the signal we see may be from effects on ORF45 transcription (**Figure 2c**). For high-throughput experiments, screening for additional phenotypes may be required to separate the multiple functions encoded by the virus.

ORF46 encodes for a uracil-DNA N-glycosylase (UNG) – a DNA repair enzyme responsible for the removal of uracil from DNA (Earl *et al.*, 2018) – and UNG homologs in EBV, human cytomegalovirus (HCMV), and vaccinia virus are all required for viral DNA replication (Stuart *et al.*, 1993; Pyles and Thompson, 1994; Ranneberg-Nilsen *et al.*, 2012; Su *et al.*, 2014). Notably, the DNA binding domain but not UNG activity is required for DNA replication in both EBV (Su *et al.*, 2014) and KSHV, suggesting that UNGs may play conserved structural rather than enzymatic roles in genome replication for multiple dsDNA viruses.

Beyond ORF46, our initial screen highlighted additional viral genes with more modest effects than the canonical late gene regulators (**Supplementary Data 4**). While their roles in late gene expression remain unvalidated, they may represent a deeper cast that either play a minor role or whose signal is limited technically. Technical limitations such as poor sgRNA performance or number may also explain the weak signal from smaller genes such as ORF30 and ORF31, two additional components of the vTA complex. An additional limitation is that less than 25% of cells detectably express the late gene reporter (**Figure S1**); while this percentage is consistent with previous reports (Brulois *et al.*, 2014; Nakajima *et al.*, 2020), cell models allowing for more efficient late gene expression or detection should further boost the dynamic range and sensitivity. While individual sgRNAs targeting the major transactivator ORF50 behaved as expected (**Figure S2b**), their positive enrichment in the screen (**Figure 1b,c**) may represent an unknown role for ORF50 or an artifact of the screen design. Additionally, positive signals were detected for KSHV genes required for latency, such as ORF73 (LANA) and the GFP/HygroR cassette (**Figure 1b,c**), but their relevancy to late gene expression is unclear given that the screen was performed under strong selection for latency maintenance.

The tiling library is highly modular, as it can be used for example with base editors to create base pair mutations, CRISPRi to induce transcriptional repression or dCas9 to block transcription factor binding, as well as in combination with other reporters of viral activity. Overall, the tiling approach is a powerful method for functional interrogation of KSHV and other DNA viruses with the added potential to highlight essential domains and unlock a wealth of information about viral biology.

**Table 1.** Number of reads and detected mutations.

|  | Latent rep 1 | Latent rep 2 | Replicated DNA<br>rep 1 | Replicated DNA<br>rep 2 | Supernatent<br>rep 1 | Supernatent<br>rep 1 |
| --- | --- | --- | --- | --- | --- | --- |
| Aligned reads | 164,795,650 | 123,368,798 | 130,643,727 | 114,902,707 | 111,795,168 | 122,456,585 |
| Total indels | 260,281 | 205,906 | 185,299 | 155,538 | 130,088 | 136,901 |
| Masked | 75,707 | 65,352 | 65,339 | 57,180 | 47,245 | 51,889 |
| Pairs | 4,338 | 3,110 | 4,055 | 2,746 | 2,063 | 2,199 |
| Duplicates | 120,312 | 88,747 | 78,278 | 62,838 | 56,532 | 57,686 |
| Simple indels | 53,310 | 42,975 | 32,835 | 28,591 | 20,383 | 21,284 |

---

**Figure 1S.**
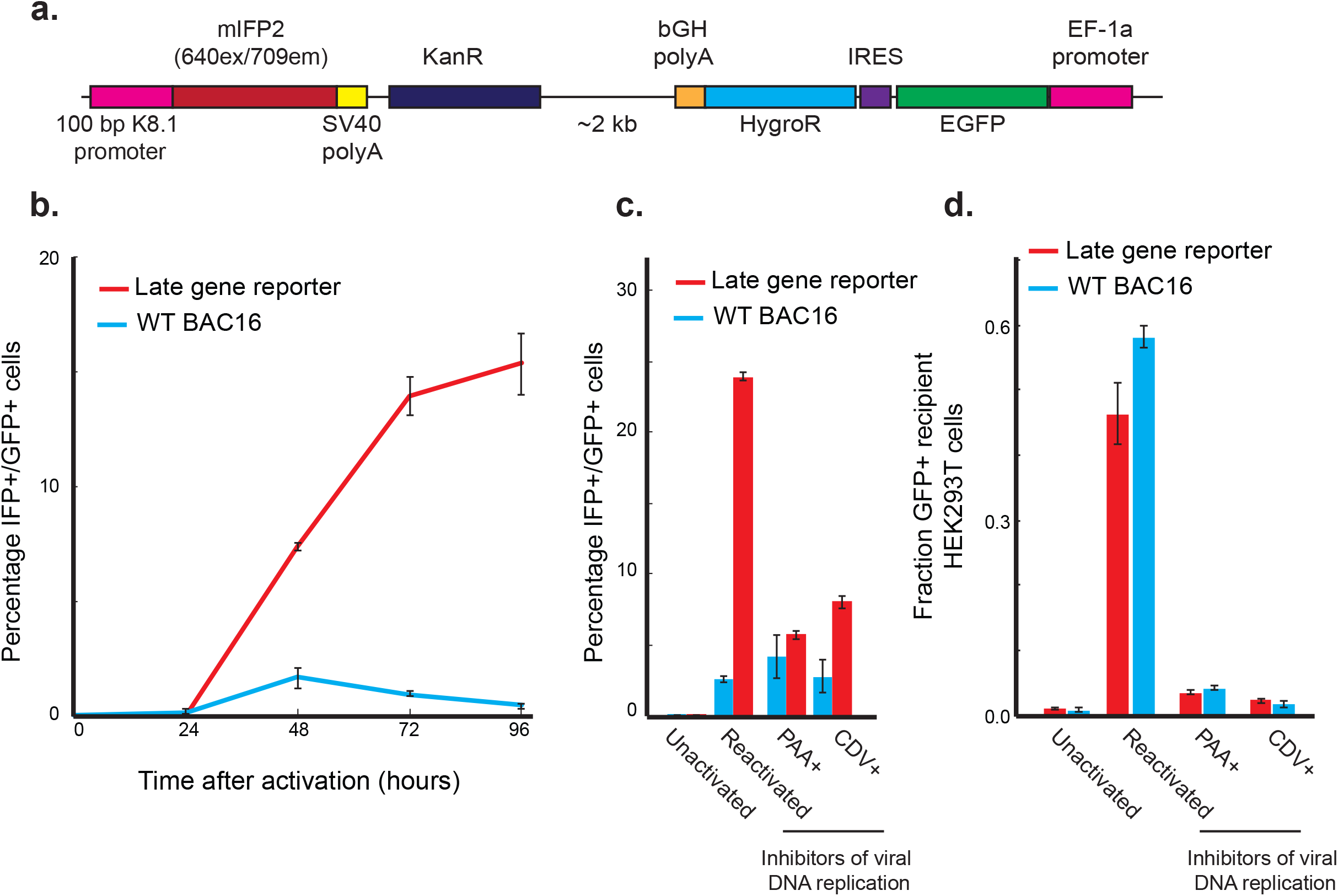
BAC16 K8.1pr-mIFP2 infected iSLK cells enables fluorescent readout of late gene activity. a) Design of a late gene reporter. A cassette expressing the far-red fluorescent protein driven by the 100 basepair promoter sequence of the KSHV late gene K8.1 was inserted upstream of the EGFP cassette of the BAC16 KSHV genome. A KanR cassette was included to allow selection in bacteria. b) The late gene reporter is expressed 48-96 hours post-reactivation. iSLK cells infected with either the wildtype BAC16 or the late-gene reporter modified BAC16 were reactivated, and reporter activity was monitored every day for 96 hours. c) Sensitivity of late gene reporter to inhibitors of viral DNA replication phosphonoacetic acid (PAA) and cidofovir (CDV). Reporter activity was monitored by flow cytometry 72 hours after reactivation. Inhibitors of viral DNA replication restricted reporter activity. d) Viral activity of late gene reporter virus after transfer of supernatant to uninfected HEK293T cells. 72 hours post-reactivation, viral supernatant was filtered and transferred to naïve HEK293T cells; infection of HEK293T cells was monitored by expression of the BAC16-encoded, constitutive EGFP. Error bars are standard error from three technical replicates.

**Figure 2S.**
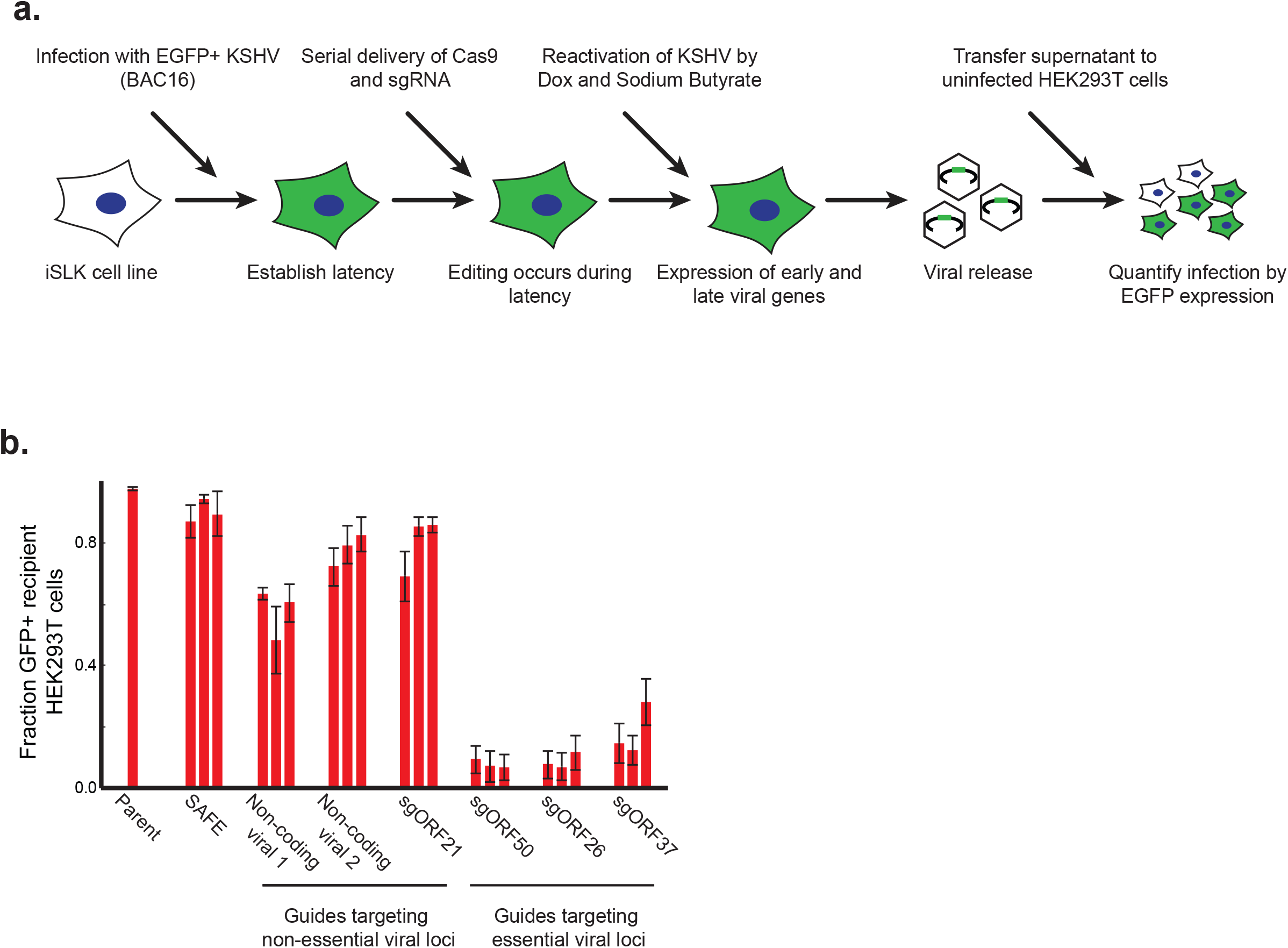
CRISPR/Cas9 enables functional targeting of viral genes. a) Delivery and editing during latent infection of iSLK cells. Cas9-blast was lentivirally delivered to BAC16 infected iSLK lines. After selection, mU6-driven sgRNAs were delivered lentivirally. Cells were maintained in a latent state for 1-2 weeks to allow sufficient time for editing before reactivation by doxycycline and sodium butyrate. b) Viral activity after supernatant transfer to uninfected HEK293T cells. SAFE denotes sgRNAs targeting the host in regions without expected function (Morgens *et al.*, 2017). Non-coding viral 1 and 2 indicates sgRNAs targeting two viral regions free of ORFs. Each bar represents an independent sgRNA, and error bars represent standard error from three independent reactivations.

**Figure 3S.**
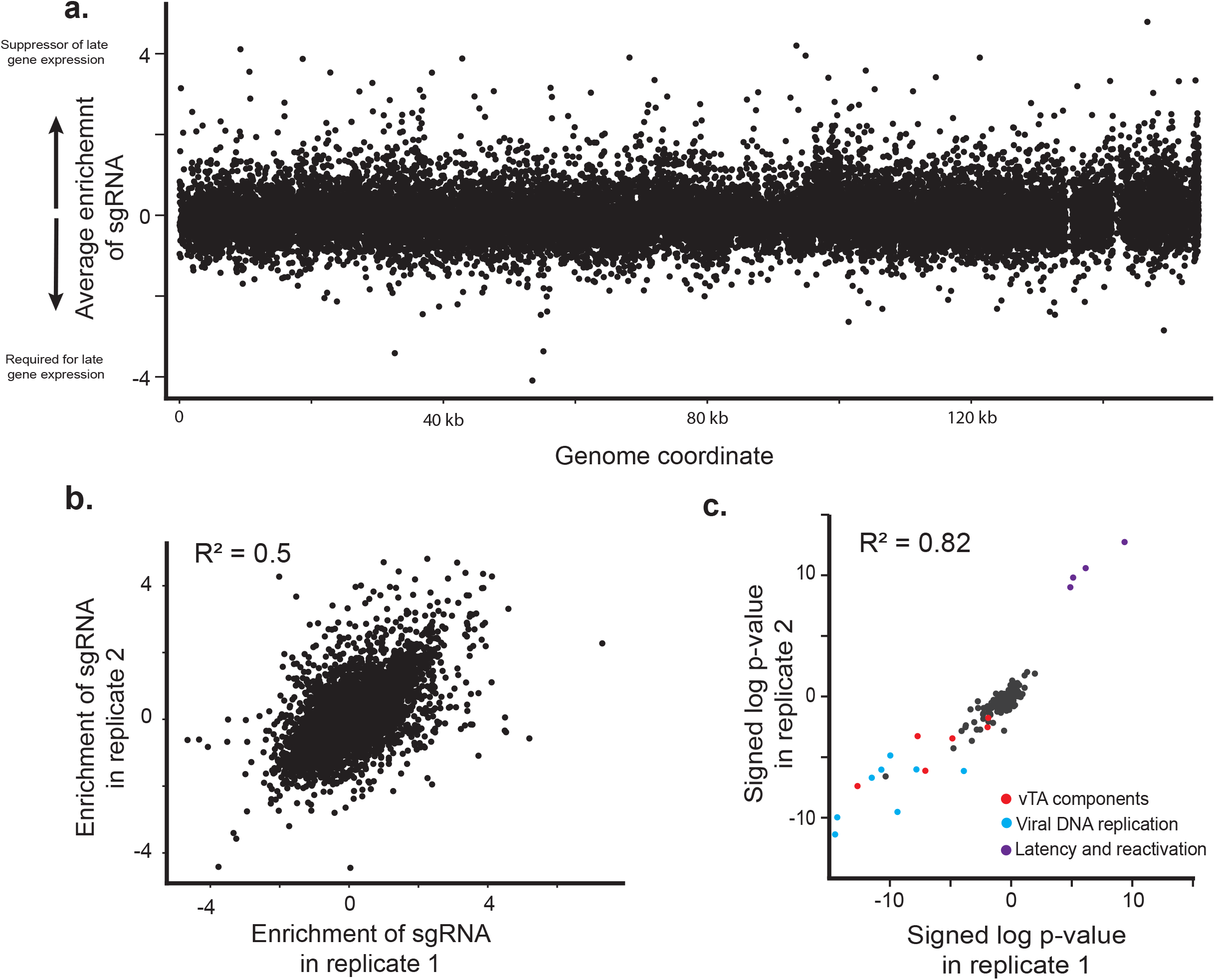
Individual results from late gene tiling screen. a) Average enrichment score from two replicates across the 154 kb genome. b) Guide-level reproducibility of sgRNA enrichments from two independent replicates. Positive value indicates the sgRNA is enriched in the late-gene expressing fraction and thus promotes late-gene expression; negative value indicates the sgRNA is depleted from the late-gene expressing fraction and thus suppresses late gene expression. c) Gene-level reproducibility of p-values from two replicates. To calculate significance, sgRNAs targeting each annotated region of the genome were grouped and a signed log Mann-Whitney p-value was calculated comparing each viral region to the negative control sgRNAs. A positive log p-value indicates this region promotes late gene expression when disrupted, and a negative log p-value indicates this region is required for expression of late genes.

**Figure 4S.**
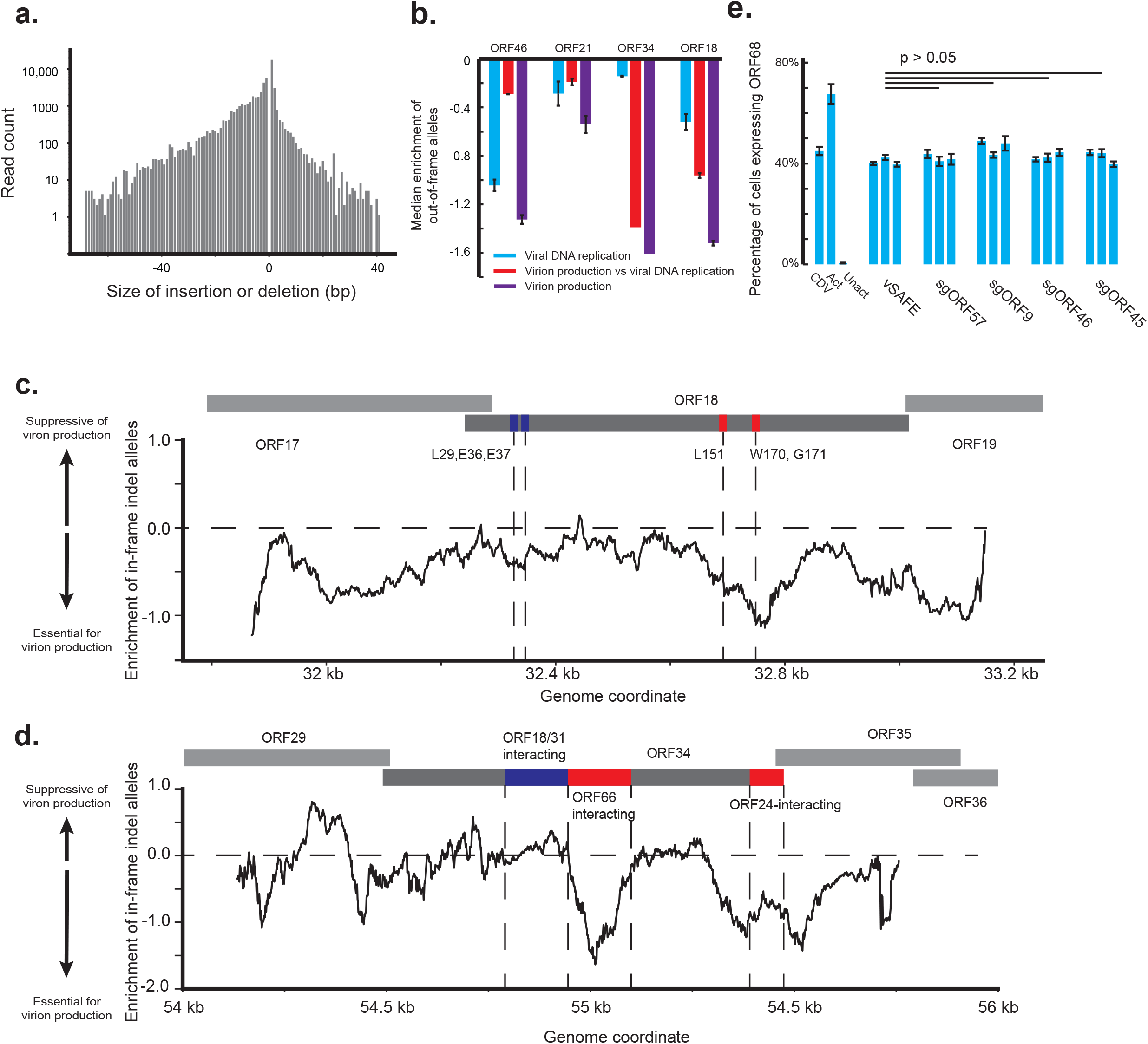
Additional results from targeted viral sequencing. a) Representative spectra of indel sizes in a single replicate of latent genome. b) Median enrichment across the coding region of the gene of out-of-frame indels in replicating cells relative to latent cells (Viral DNA replication), the supernatant relative to latent cells (Virion production), or supernatant relative to replicating cells (Viral DNA replication vs Virion production). Error bars are standard error from two replicates. One replicate of ORF34 supernatant sample was excluded due to uneven coverage. c,d) Smoothed signal from in-frame mutations across the c) ORF18 and d) ORF34 loci in comparison between replicating and supernatant genomes. Functional domain labeling for ORF18 (Castañeda and Glaunsinger, 2019) and ORF34 (Nishimura *et al.*, 2017). e) Reactivation and early gene expression was measured using a KSHV virus containing a HaloTag fusion to a viral early gene, ORF68. vSAFE indicates sgRNAs targeting a viral region free of ORFs. Each bar is an independent sgRNA, and error bars are standard error from four independent reactivations. P values were calculated using a single-tailed, equal variance Student’s t-test, with the least significant value used when compared to each individual vSAFE sgRNA. NS indicates non-significance (P>0.05).

**Supplementary Table 1.** Number of reads and detected mutations.

**Supplementary Data 1.** List of sgRNA sequences in KSHV tiling library

**Supplementary Data 2.** sgRNA counts from late gene screen

**Supplementary Data 3.** Genome reference and annotation

**Supplementary Data 4.** Results of late gene screen

**Supplementary Data 5.** Detected mutations for viral sequencing

**Supplementary Data 6.** Frequencies for viral sequencing

**Supplementary Data 7.** Sequence of sgRNAs and primers used

## Methods

### Plasmids and oligos

Sequences of primers and sgRNAs used are listed in **Supplementary Data 7.** pMD2.G (Addgene plasmid # 12259), pMDLg/pRRE (Addgene plasmid # 12251) and pRSV-Rev (Addgene plasmid # 12253) were gifts from Didier Trono. pMCB320 was a gift from Michael Bassik (Addgene plasmid # 89359). lentiCas9-Blast was a gift from Feng Zhang (Addgene plasmid # 52962).

### Cell culture

HEK293T and iSLK cells were grown in DMEM (Gibco, + glut, + glucose, -pyruvate) with 10% FBS (Peak Serum), pen-strep (Gibco; 10,1000 I.U./mL), and additional 2 mM glutamine (Gibco) or 1X Glutamax (Gibco). Cells were maintained at 37°C degrees and 5% CO_2_ in a humidity-controlled incubator. iSLK cells were maintained in 1 μg/mL puromycin and 50 μg/mL G418, with infected iSLK cells additionally maintained in 125-200 μg/mL Hygromycin. Cas9+ cells were maintained in 10 μg/mL blasticidin. 0.05% Trypsin (Gibco) was used to passage cells.

### Creation and characterization of late gene reporter cell line

A late gene reporter was designed using 100 bp upstream of the K8.1 ORF driving a mIFP2 fluorescent cassette. 150 bp of homology to the BAC region of BAC16 was included along with a KAN resistance cassette to allow the insertion into BAC16 genome (Brulois *et al.*, 2012). The viral genome was then purified using the Macherey-Nagel NucleoBond BAC 100 kit and transfected into HEK293T cells using PolyJet (SignaGen). Transfected cells were then cocultured with uninfected iSLK cells and reactivated using 0.33 mM sodium butyrate and 20 ng/mL PMA. Infected iSLK cells were then selected using 1 μg/mL puromycin, 50 μg/mL G418, and 200 μg/mL hygromycin.

To test the kinetics of late-gene reporter expression, iSLK-BAC16-K8.1pr-mIFP2 cells were reactivated using 1 mM sodium butyrate and 5 μg/mL doxycycline, and timepoints were collected every 24 hours. Cells were then fixed in 4% PFA and quantified using flow cytometry (BD LSRFortessa). To test the sensitivity of the late gene reporter-infected lines, cells were reactivated as above and additionally treated with viral polymerase inhibitors 500 μM PAA or 100 μM CDV; late-gene reporter activity of fixed cells was quantified by flow cytometry 72 hours post reactivation (BD LSRFortessa). To test the viral activity of these cells, supernatant from reactivated samples was 45 um filtered and added to uninfected HEK293T cells. After 24 hours, HEK293T cells were fixed, and GFP was quantified by flow cytometry (BD Accuri C6 Plus). Replicates were independent reactivations on separate days.

### Creation and characterization of Cas9 positive line

To create a Cas9 positive iSLK-BAC16-K8.1pr-mIFP2 line, Cas9-blast (Addgene #52962) was inserted lentivirally: Cas9-blastR and 3^rd^ generation lentiviral components were transfected into HEK293T cells using PEI (Polysciences). Supernatant was harvested, 45um filtered, and added to iSLK-BAC16-K8.1pr-mIFP2 line for three days. Cells were then selected using 1 μg/mL blasticidin, 1 μg/mL puromycin, 200 μg/mL hygromycin, and 50 μg/mL G418.

To demonstrate the editing activity, three independent sgRNAs were designed targeting two noncoding regions of the BAC and four viral genes. sgRNAs were designed using IDT Custom Alt-R designer. As controls, three safe-targeting sgRNAs were used that target the host in predicted non-functional regions (Morgens *et al.*, 2017). sgRNAs were cloned into a mU6-driven guide expression plasmid (Addgene #89359) and delivered lentivirally at high MOI. After 10 days, cells were reactivated by 5 μg/mL doxycycline and 1 mM sodium butyrate, and viral supernatant was collected and 45um filtered after 72 hours. Supernatant was transferred to uninfected HEK293T cells, and viral infectivity was measured using GFP by flow cytometry (BD Accuri C6 Plus). Replicates were independent reactivations on separate days.

### Design of KSHV tiling library

A KSHV tiling library was designed using custom python scripts. Each NGG PAM was identified in the reference genome (Brulois *et al.*, 2012) (**Supplementary Data 1, 3**) along with the corresponding 19 bp guide. Only one copy of sgRNAs that targeted the KSHV genome more than once was included. The library was cloned using a modified protocol from Deans et al 2016(Deans *et al.*, 2016). sgRNAs were synthesized along with primer binding sites and BstXI/BlpI restriction sites using Twist Biosciences. The library was amplified using corresponding primers for 10 cycles. Amplified library was PCR purified (Qiagen MinElute) and restricted using BstXI and BlpI. 34 bp fragment was purified in water using a native 20% PAGE gel and Spin-X centrifuge tube filters (Costar). Fragment was isoproponal precipitated and ligated into a sgRNA expression plasmid (Addgene #89359) using T4 ligase overnight at 16 degrees and electroporated into Lucigen Endura cells (1800V, 600 ohm, 10μF, 1mm) before plating on 4 assay plates (reduced 75% CARB). Colonies were grown at 37 degrees C overnight, pooled, and maxi prepped (Macherey-Nagel).

### Screen and analysis of late gene screen

Cas9+ iSLK-BAC16-K8.1pr-mIFP2 cells were infected lentivirally with the KSHV tiling library. After 10 days, cells were reactivated using 1 mM sodium butyrate and 5 μg/mL doxycycline. Two days post reactivation, cells were trypsinized, washed 2x with dPBS, and fixed in 4% PFA for 10 minutes before being washed 2x in dPBS again. Cells were then sorted based on mIFP2 expression using a BD Aria, sorting for top 25% expression vs bottom 50% expression (BD Aria). Sorted cells were then unfixed overnight in 50 μg/mL proteinase K (Promega) plus 150 mM NaCl at 65°C. Genomic DNA was extracted using the Qiagen blood mini kit. As described in Deans et al 2016 (Deans *et al.*, 2016), the sgRNA locus was amplified using primers and Herculase II (Agilent) with 20 cycles. Nextera-index adapters were then ligated by PCR using 18 cycles. ~285 bp product was gel extracted and quantified with Qubit and an Agilent Bioanalyzer before sequencing on a HiSeq 4000.

Average enrichment of sgRNAs from two replicates were calculated comparing the late-gene high and late gene low fractions using a median normalized log ratio of fraction of counts as previously described (Kampmann, Bassik and Weissman, 2013). sgRNAs that targeted the viral genome at multiple loci were removed from the analysis. Each region of the viral genome was split using previous annotations (**Supplementary Data 3**), and the median enrichment of all sgRNAs targeting each region was compared to all negative control sgRNAs using a Mann-Whitney U test.

### Screen validation, early gene expression, and viral DNA replication

Three sgRNAs from the screen were selected, cloned as above, lentivirally delivered to late gene expression reporter cells, and assayed for viral activity and late gene expression activity as above. sgRNAs were also delivered to Cas9+ iSLK cells infected with a modified BAC16 virus containing an N-terminal HaloTag fusion to the early viral gene ORF68. 24 hours post reactivation with 1 mM sodium butyrate and 5 μg/mL doxycycline, cells were treated with 20 nM HaloTag JF647 (Promega) and ORF68 levels were quantified by flow cytometry (Accuri C6 plus). To quantitate viral DNA replication, 20 μM EdU was added to the media 48 hours post reactivation for two hours. Cells were then trypsinized and fixed using 4% PFA. EdU was labeled with Cy5 using the Click-IT flow cytometry kit (Invitrogen) and quantified by flow cytometry (BD Accuri C6 Plus). Using unreactivated cells as a control, little host DNA replication was observed in reactivated cells. EdU levels below those observed in unreactivated cells but above background were quantified as viral DNA replication. To quantitate the amount of viral DNA, 48 hours after reactivation, DNA was extracted using DNA QuickExtract (Lucigen), and qPCR was performed using iTaq Universal SYBR Green (Bio-rad) amplifying the ORF59 promoter region of KSHV (QuantStudio 3). Viral quantities were calculated by standard curve. Replicates were independent reactivations on separate days.

### Design and analysis of targeted viral sequencing

Cas9+ iSLK-BAC16-K8.1pr-mIFP2 cells containing the KSHV tiling library were reactivated using 1 mM sodium butyrate and 5 μg/mL doxycycline. Genomic DNA was recovered from latent cells prior to reactivation and 48 hours post reactivation using a QIAamp DNA blood Mini kit (Qiagen). To collect viral supernatant, media was replaced 48 hours post reactivation and collected 72 hours post reactivation. Viral supernatant was then mixed with appropriate volume of Qiagen AL buffer and Qiagen proteinase. Sample was then spun over a single QIAamp blood mini column, washed, and eluted as the manual indicates.

PCR was performed from each DNA sample for four regions (ORF21, ORF34, ORF46, and ORF18) using Herculase II (Agilent). Each region was then PCR purified (GeneJet; Thermo) and pooled in equal nanogram amounts. 500 ng of each pooled sample was then prepped using Illumina DNA prep and sequenced on a single lane of SP 150PE NovaSeq 6000.

Reads were trimmed using cutadapt (Martin, 2011), and aligned to the BAC16 genome using bwa mem(Li and Durbin, 2009). Indel counts were then extracted from CIGAR codes, with identical CIGAR codes excluded as duplicates. Complex codes not corresponding to simple indels were excluded. Deletions and insertions at the same position and same size observed in a control sample amplified from purified BAC were excluded as artifacts of library prep (**Supplementary Data 5, Supplementary Table 1**). Insertions or deletions whose size were divisible by three were classified as in-frame mutations and all others were classified as out-of-frame mutations. Mutation counts and coverage were calculated as the sum of two replicates for each condition. Basepairs with less than 10^5 coverage were excluded, and a single read was added to each basepair to prevent division by zero. Mutation frequencies across each 100 bp were averaged and a log enrichment value was calculated between either the latent sample and the 48 hour sample, or the 48 hour sample and the supernatant sample. For each comparison, the difference in log enrichment between in-frame mutations and out-of-frame mutations was calculated and used as the relevant signal.

### ORF46 BAC mutagenesis

ORF46 DNA binding (H210L) or catalytic (Q87L, D88N) mutants were introduced into BAC16 using red recombination (Brulois *et al.*, 2012). Briefly, homology arms were used to insert mutations along with a KAN resistance cassette into the endogenous ORF46 locus. The KAN cassette was removed by recombination, and the scarless edit was confirmed by Sanger sequencing. Viral genomes were then purified using the Macherey-Nagel NucleoBond BAC 100 kit and transfected into HEK293T cells using PolyJet (SignaGen). Transfected cells were then cocultured with uninfected iSLK cells and reactivated using 0.33 mM sodium butyrate and 20 ng/mL PMA. Infected iSLK cells were then selected using 1 μg/mL puromycin, 50 μg/mL G418, and 125 μg/mL hygromycin.

iSLK cells infected with the mutant BAC16 viruses were then reactivated using 5 μg/mL doxycycline and 1 mM sodium butyrate. After 72 hours, supernatant was 45um filtered and transferred to uninfected HEK293T cells. 24 hours later, infection was monitored by EGFP expression using flow cytometry (BD Accuri C6 plus). Replicates were independent reactivations from a single experiment.

## Author Contributions

D.W.M. and B.G. conceived and designed the study and wrote the manuscript. D.W.M. designed the library and performed screens and analyses. D.W.M. and D.N. collated annotations. D.N. designed and validated EdU viral detection assay. A.L.D. designed and created ORF68 HaloTag iSLK line. D.W.M. and K.Y. performed mutant follow-up assays. All authors reviewed and approved the manuscript.

## Acknowledgements

We thank E. Hartenian, X.J. Mao, L. Coscoy, and members of the Glaunsinger and Coscoy labs for helpful discussion. We thank C.K. Tsui for their support and review of the manuscript. Flow cytometry and FACS were conducted at the CRL Flow Cytometry Facility. We thank Hector Nolla and Alma Valeros of the UC Berkeley Cancer Research Laboratory Flow Cytometry Facility for training and expertise. This work used the Vincent J. Coates Genomics Sequencing Laboratory at UC Berkeley, supported by NIH S10 OD018174 Instrumentation Grant. D.W.M. is a Howard Hughes Medical Institute Awardee of the Life Sciences Research Foundation. A.L.D. is the Rhee Family Fellow of the Damon Runyon Cancer Research Foundation (DRG-2349-18). Work was supported by a Research for Innovation on Delivery of Editing Reagents award from the Innovative Genomics Institute and the NIAID (2R01AI122528-06 to B.G.). B.G. is an investigator for the Howard Hughes Medical Institute.

